# Mind your gaps: Overlooking assembly gaps confounds statistical testing in genome analysis

**DOI:** 10.1101/252973

**Authors:** Diana Domanska, Chakravarthi Kanduri, Boris Simovski, Geir Kjetil Sandve

**Affiliations:** Department of Informatics, University of Oslo, Oslo, Norway; K. G. Jebsen Coeliac Disease Research Centre, Oslo, Norway.

**Keywords:** assembly gaps, reference genome, statistical genome analysis, co-localization analysis, co-occurrence analysis, region set enrichment analysis, genomic overlap analysis, ChIP-seq, RNAseq; GWAS, genomic regions, genomic intervals, shuffling, regulatory regions

## Abstract

**Background:** The difficulties associated with sequencing and assembling some regions of the DNA sequence result in gaps in the reference genomes that are typically represented as stretches of Ns. Although the presence of assembly gaps causes a slight reduction in the mapping rate in many experimental settings, that does not invalidate the typical statistical testing comparing read count distributions across experimental conditions. However, we hypothesize that not handling assembly gaps in the null model may confound statistical testing of co-localization of genomic features.

**Results:** First, we performed a series of explorative analyses to understand whether and how the public genomic tracks intersect the assembly gaps track (hg19). The findings rightly confirm that the genomic regions in public genomic tracks intersect very little with assembly gaps and the intersection was observed only at the beginning and end regions of the assembly gaps rather than covering the whole gap sizes. Further, we simulated a set of query and reference genomic tracks in a way that nullified any dependence between them to test our hypothesis that not avoiding assembly gaps in the null model would result in spurious inflation of statistical significance. We then contrasted the distributions of test statistics and p-values of Monte Carlo simulation-based permutation tests that either avoided or not avoided assembly gaps in the null model when testing for significant co-localization between a pair of query and reference tracks. We observed that the statistical tests that did not account for the assembly gaps in the null model resulted in a distribution of the test statistic that is shifted to the right and a distribu tion of p-values that is shifted to the left (leading to inflated significance).

**Conclusion:** Our results shows that not accounting for assembly gaps in statistical testing of co-localization analysis may lead to false positives and over-optimistic findings.

## Content

Text and results for this section, as per the individual journal’s instructions for authors.

## Background

Genome biology research relies on reference genomes to a large extent to map the high-throughput sequencing reads against known functional annotations [1]. The reference genome thus serves as a central entity that interlinks various genomic features [2]. Therefore, our current understanding of the genomes is greatly influenced by the completeness of the reference genomes [3]. However, because of the challenges associated with cloning and mapping certain highly repetitive and complex regions, the physical maps of the reference genomes of many species currently contain long-stretches of gaps [4]. For instance, a considerable proportion of the human genomic sequence (between 5-10%) remains poorly characterized to date. In the latest human genome build, hg38, around 200 Mbp mainly from centromeres and acrocentric short arms, and around 30 Mbp of interstitial gaps mostly in euchromatic sequences are currently uncharacterized [5]. The genome assemblies of non-model organisms contain a higher gap proportion than humans and model organisms (e.g., 6% gap bases in the genome of giant panda). [6, 7].

High-throughput sequencing reads are not expected to map against the assembly gap regions of the reference genome because of the lack of reference sequences. In many experimental settings, the little reduction in mapping rate due to the presence of assembly gaps is not a major constraint (owing to the relatively smaller size of the gap regions compared to the whole genome). However, we hypothesise that the presence of assembly gaps and not handling them in the null model can introduce a bias and lead to over-optimistic findings in statistical hypothesis testing of co-localization of genomic tracks.

The completion of the human genome sequencing and the subsequent improvements in the sequencing technologies have allowed a deeper understanding of the genomic sequences and various genomic features. Over the past decade, understanding the interplay of genomic features has become a crucial direction in biomedical research, especially in understanding the molecular genetic background of diseases and traits. One important way to understand the interplay of various genomic features is to quantitate the overlap or co-localization of genomic features and test the statistical significance of the observed co-localization. Monte Carlo (MC) simulations-based permutation tests are a preferred way to perform the statistical hypothesis testing, where one could avoid over-simplistic and over-optimistic null models that are typical of simplistic analytical solutions like Fisher’s exact test [8, 9]. Irrespective of the choice of the statistical testing approach, one should essentially avoid the assembly gap regions in the null model (here it is noteworthy that all the statistical tests assume a null model; [8, 9]). General purpose genome arithmetic tools that can shuffle genomic tracks typically provide a functionality to restrict the shuffling to certain regions, where one could restrict the shuffling in assembly gaps [10]. However, performing a MC simulations-based permutation test on such collection of shuffled tracks involves ad hoc choices confounded by simplistic null models and thus not recommended [8, 9]. On the other hand, simplistic analytical solutions that for instance use a Fisher’s exact test cannot account for assembly gaps as Fisher’s exact test only preserves the total number of bases or intervals in the null model, but not other characteristics of the real data and therefore not a recommended choice either. Although some co-localization analysis tools based on MC simulations allow the shuffling of genomic elements restricted by some regions (where one can essentially avoid assembly gaps), the user needs to be aware of the magnitude of potential bias and the possibility of over-optimistic findings because of not avoiding assembly gaps. Here, we demonstrate the potential bias introduced when assembly gaps are not appropriately handled, and show that this can often lead to over-optimistic or false-positive findings.

## Results and Discussion

### Overlap of public genomic tracks with genome assembly gaps

We performed a series of explorative analyses to understand the nature and extent of the intersection of genomic tracks with genome assembly gaps. For this, we downloaded large collections of public genomic tracks (hg19) categorized by diverse experimental assays, including tracks of histone modifications, DNase I hypersensitive sites, and transcription factor binding sites in the K562 cell line in table 1. We then intersected the genomic tracks with assembly gaps of hg19. Overall, we observed a trend of a very modest overlap with the assembly gaps and the overlap was unsurprisingly localized to the beginning and end portions of the gaps, rather than crossing over the middle portion of the gaps in table 2. Assembly gap regions that overlapped the most with the public genomic tracks are on the chr4:40296396-40297096, chr7:139379377-139404377 followed by chr3:194041961-194047251. These findings rightly confirm that the sequencing reads do not map to the assembly gaps on hg19. This is not surprising, because unlike hg38, which contains sequence models for a large portion of the gaps, hg19 rather contains long-stretches of Ns in most of the gap regions, thus excluding the possibility of any read mapping.

**Table 1:** Track collection and aim of analysis.

| Tracks collection | Number of tracks | Source of Genomic tracks | Purpose |
| --- | --- | --- | --- |
| Histone modifications | 477 | ENCODE | Investigating the nature and extent of overlap of public genomic tracks with genome assembly gaps |
| DNase I hypersensitive sites | 838 |  |  |
| Transcription factor binding sites (K562) | 568 |  |  |
| Randomly selected tracks from the above collections | 100 | ENCODE | Understand the impact of not avoiding assembly gaps on the conclusions of statistical colocalization analysis |

**Table 2:** Descriptive statistics of the intersection of public genomic tracks (counted relatively to the gaps size) with genome assembly gaps (hg19).

| Tracks collection | Avg size of genomic intervals (bp) | Avg overlap with gaps (bp) | Avg overlap with the start region of gaps (bp) | Avg overlap with the end region of gaps (bp) | Avg overlap with the middle portion of gaps (bp) | Overlap hotspot | Overlap hotspot value (bp) |
| --- | --- | --- | --- | --- | --- | --- | --- |
| Histone modifications | 2.26e-13 | 0.0014 | 0.0029 | 0.0023 | 0.0011 | chr4:40296396-40297096 | 0.0591 |
| TFBS in K562 | 1.28e-12 | 0.0046 | 0.0059 | 0.0053 | 0.0046 | chr7:139379377-139404377 | 0.0448 |
| DNase I hypersensitive sites | 3.68e-15 | 4.49e-06 | 3.28e-05 | 1.21e-05 | 0 | chr3:194041961-194047251 | 0.0007 |

We further checked whether the amount of overlap of genomic tracks with assembly gaps is size-dependent. We observed that the overlap of public genomic tracks with assembly gaps increased with an increase in the average segment length of the tracks (Supplementary materials, fig. 3).

### The impact of not avoiding the assembly gaps on the findings of statistical testing in co-localization analysis

To understand whether avoiding/not avoiding the assembly gap regions in the null model would have an impact on the conclusions of statistical co-localization analysis, we contrasted the findings of MC-based permutation tests that either avoided or not avoided assembly gaps in the null model. For this, we first tested the significance of pairwise overlap of 100 pairs of genomic tracks by not accounting for assembly gaps. Each of the pair of genomic tracks is comprised of a real track obtained from public repositories, while the other track was simulated to match the real track in its nature of not mapping to the assembly gap regions. By simulating genomic tracks in this fashion, one from the outset can be sure that there is no dependence (association) between the real and simulated tracks except their shared avoidance of assembly gap regions (H0 is always true). However, when not accounting for assembly gaps in the statistical tests, the distribution of p-values of co-localization analysis is strongly shifted to the left, with H0 being falsely rejected after multiple testing correction (FDR< 0:05) for 87 out of 100 tests (counted for tracks of histone modifications 1b). This shows that ignoring gaps in the null model introduces a substantial risk of false discoveries. The bias of the analysis is also evident from a comparison of the distribution of observed test statistic values (number of bases overlapping) and the average values for the test statistic under the null model, where the average observed test statistic is higher than the average test statistic of the null model when ignoring assembly gaps (Figs. 1a and 1b and Supplementary materials, figs. 4a, 4b, 5a, 5b). This trend of decreased overlap under the null model is not surprising since the null model is distributing the genomic elements uniformly without any restrictions, whereas both observed genomic tracks share an avoidance of assembly gap regions. However, the extent to which this moderate bias in overlap leads to a large number of falsely rejected null hypotheses even after multiple testing correction is notable. The severity of false rejections will depend on the statistical power of the analysis - since the assembly-gap-ignorant null hypotheses are not technically true, they will all be rejected given enough data (also after multiple testing correction). Thus, the issue of assembly gap bias may be highly problematic for datasets with many genomic regions, while it will be affecting conclusions to a lesser degree for datasets containing fewer genomic regions (see Fig. 2 and Supplementary materials, fig. 6).

**Figure 1:**
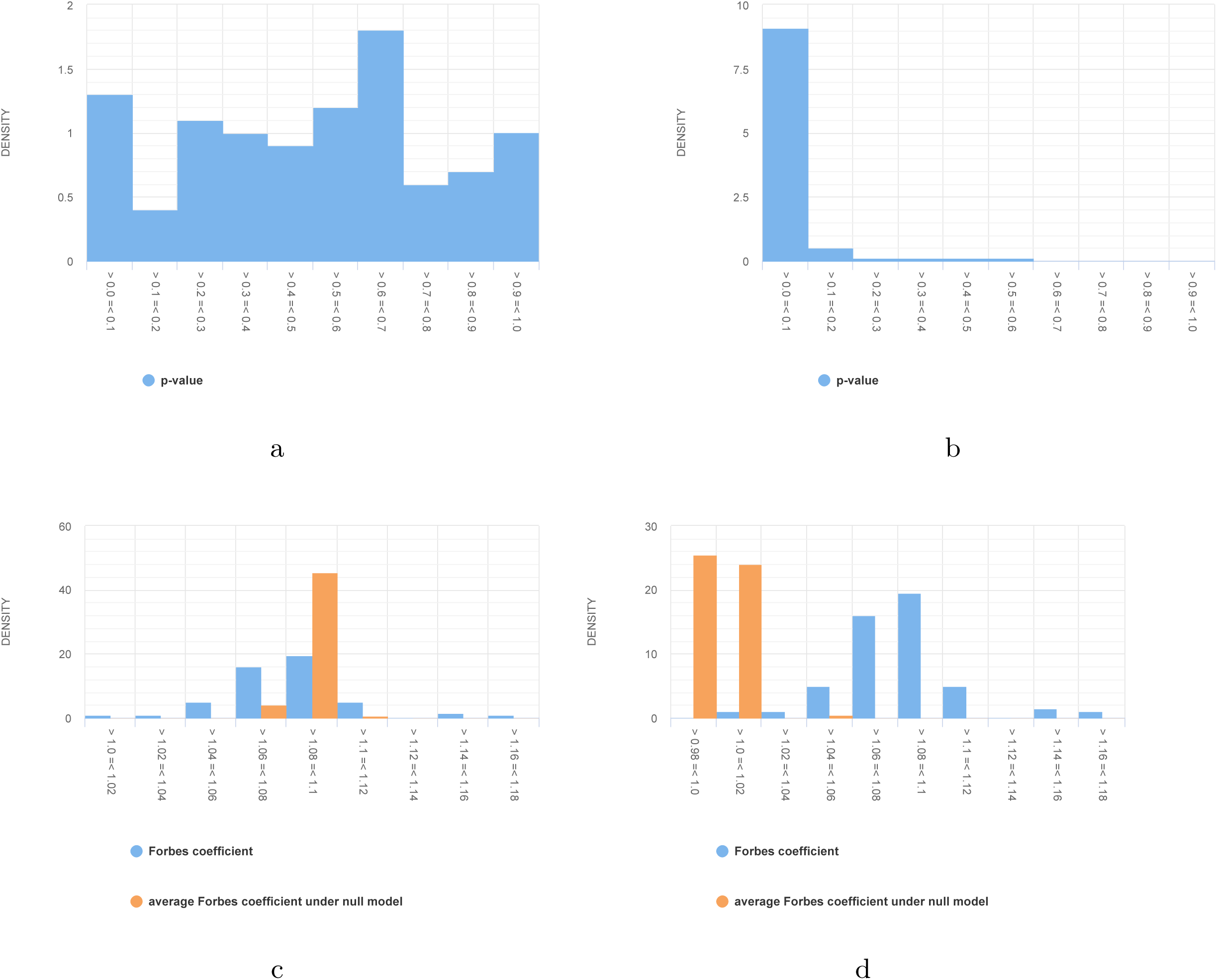
Distribution of the test statistic and p-values of co-localization analysis for a collection of 477 genomic tracks with 2113.82 bp average segment length for histone modifications. [(a) and (b) shows the distribution of p-values of the co-localization analysis with (left) and without (right) exclusion of assembly gap regions under the null model. (c) and (d) shows the observed test statistic and the average test statistic of the same tracks with (left) and without (right) exclusion of assembly gap regions under the null model. Note: Both values are higher than 1 because the computations were performed relative to the whole genome size.]

**Figure 2:**
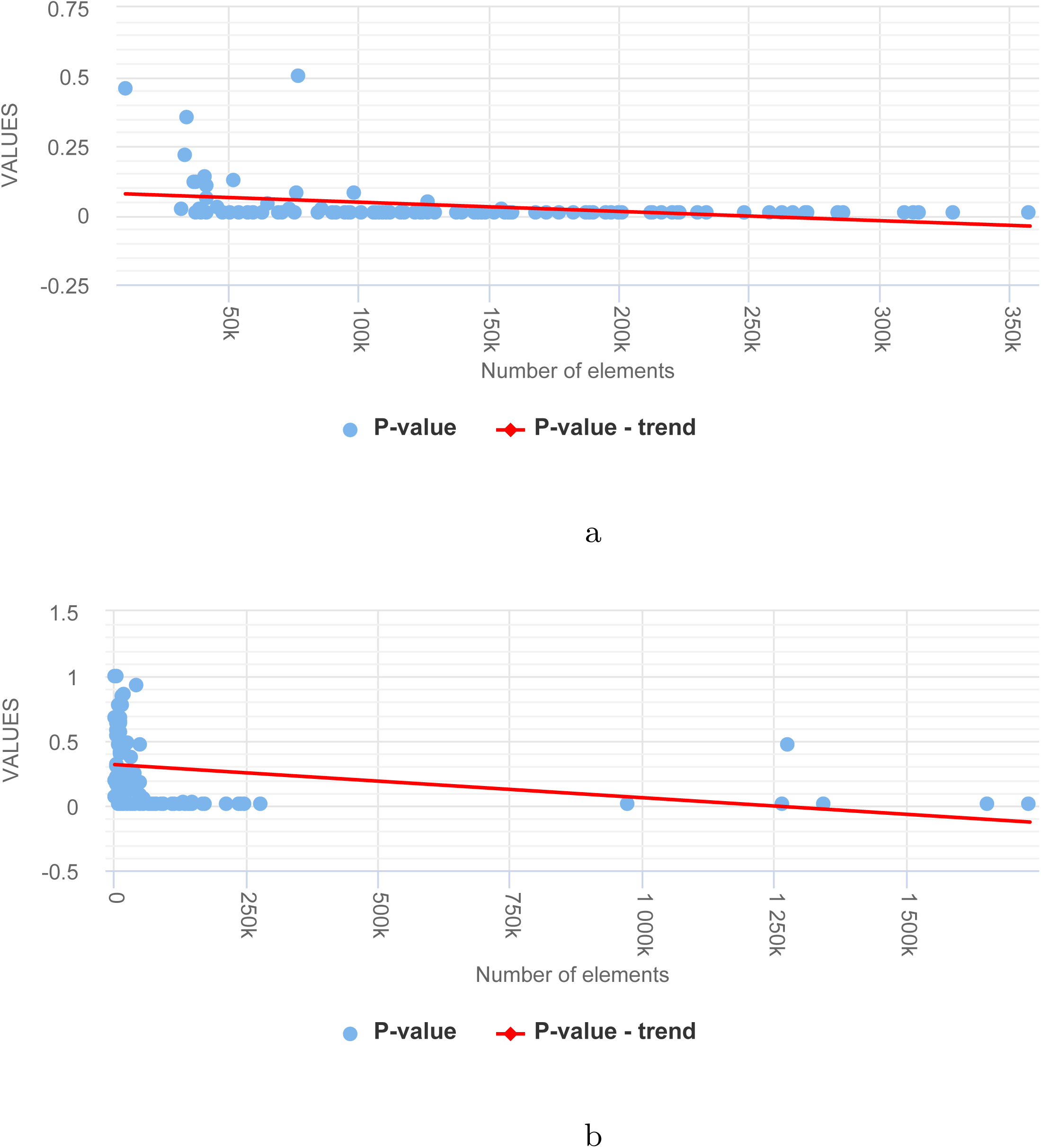
Relation between the p-values of co-localization analysis for a collection of N genomic tracks and the number of elements within each track (a) for histone modifications (N=477) (b) for TFBS in K562 (N=568).

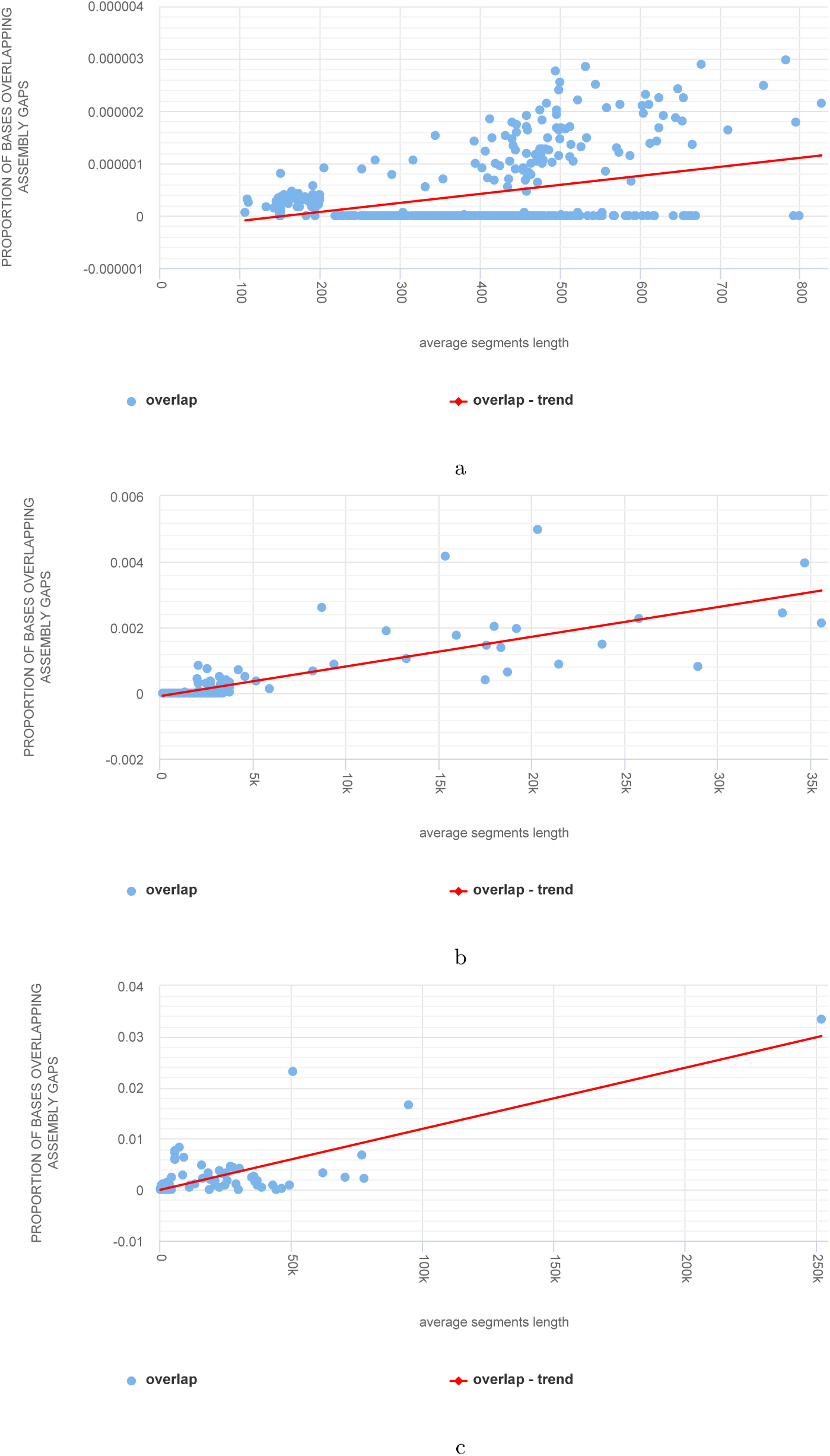

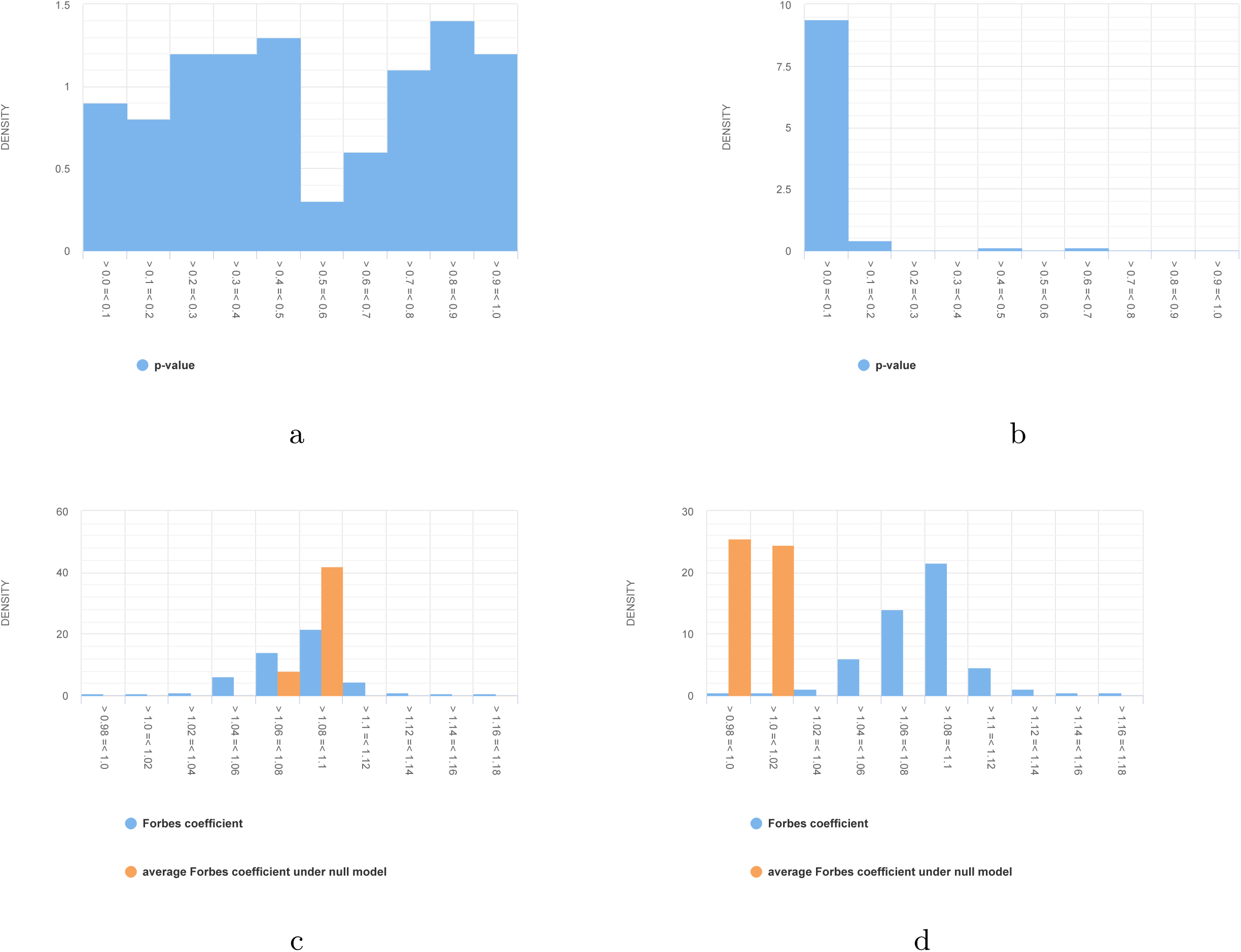

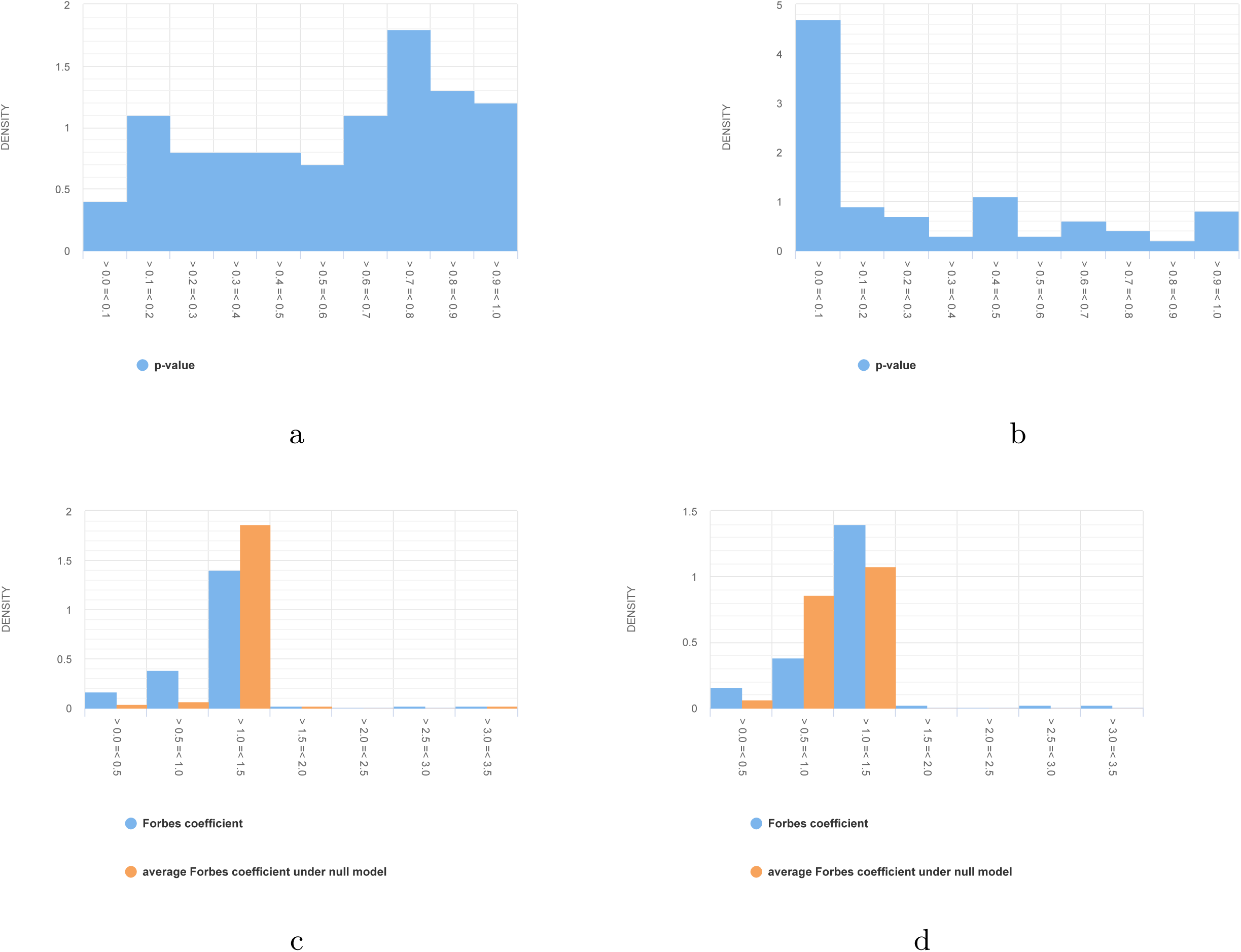

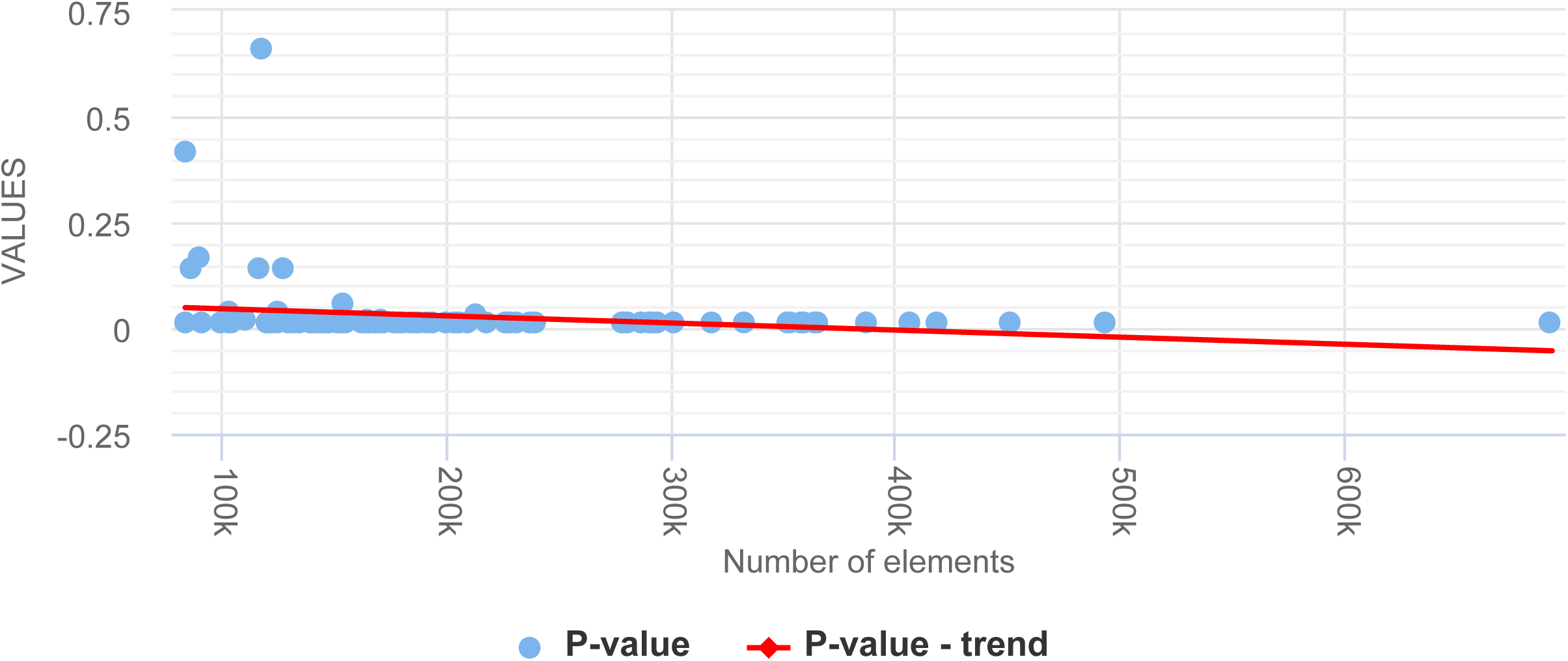

Furthermore, we repeated the statistical testing on the same dataset of 100 pairs of genomic tracks, this time by avoiding the assembly gaps in the MC simulations null model. Accounting for the assembly gaps resulted in uniformly distributed p-values with no obvious shift in either direction (Figs. 1a and 1b and Supplementary materials, figs. 4a, 4b, 5a, 5b). The comparison of the distributions of the observed test statistic and the average test statistic under the null model also substantiated that the null model is bias-free (Figs. 1c, 1d and Supplementary materials, figs. 4c, 4d, 5c, 5d).

## Methods

### Datasets

Throughout this study, we used large collections of genomic tracks that were either obtained from public repositories or simulated using a shuffling algorithm. Table 1 shows the genomic track collections used in this study. To investigate the nature and extent of overlap of genomic tracks with genome assembly gaps, we downloaded large collections of genomic tracks (hg19) from ENCODE database. The downloaded genomic tracks are categorized by diverse experimental assays including tracks of histone modifications, DNase I hypersensitive sites, and transcription factor binding sites in K562 cell line.

To understand the impact of either avoiding or not avoiding assembly gaps on the findings of statistical testing of co-localization analysis, we downloaded genomic tracks corresponding to histone modification sites (hg19) from ENCODE database and randomly retained 100 tracks for the statistical analyses. For performing pairwise co-localization analysis (based on MC simulations) on the retained tracks of histone modifications, we simulated 100 synthetic tracks that match the histone modification tracks in terms of the number of genomic regions, the average length of the genomic regions, and the distribution of genomic regions across the chromosomal arms. The synthetic tracks are also deliberately simulated in such a way that they also avoid the genome assembly gaps, mimicking the typical nature of public genomic tracks. The synthetic tracks are simulated using a standard shuffling algorithm that distributes genomic elements uniformly across the genome by avoiding assembly gap regions. For this analysis, we restricted the dataset size to 100 tracks because of the computational time of the MC simulations.

### Tools

All plots and the information necessary for their reproduction can be found at https://hyperbrowser.uio.no/assemblygaps. All results can be reproduced using the redo-functionality provided by the underlying Galaxy system (https://hyperbrowser.uio.no/assemblygaps/u/hb-superuser/p/assembly-gaps).

## Conclusions

Testing the significance of co-localization of genomic features is a common analysis approach in biomedical research. One common characteristic of genomic features assayed by current generation sequencing platforms is that the sequencing reads do not map to the gap regions of the reference genome, as also substantiated in this study. This selective depletion of reads in certain regions of the genome means that any subsequent statistical analysis should also carefully recapitulate this technical feature to avoid any potential bias. The same point holds for other scenarios in which the tracks to be analyzed share a restriction to certain parts of the genome. This could, for instance, be datasets that are restricted to coordinates in transcribed regions or regions included on a custom chip or microarray. In such situations, data should in the null model be restricted to occur in the parts of the genome where observed data could technically occur. Note that although one should always aim for null models that realistically represents technical restrictions for the real data, the bias discussed in the present paper only applies when the datasets to be analyzed share a set of excluded regions (if data is unobtainable in certain regions only for one of the tracks, it will not lead to a systematic bias as described here). This study demonstrates that overlooking assembly gap regions can confound the findings of statistical co-localization analysis and spuriously inflate the statistical significance.

## Abbreviations

MC: Monte Carlo

## Declarations

### Ethics approval and consent to participate

Not applicable

### Consent for publication

Not applicable

### Availability of data and materials

All plots and the information necessary for their reproduction can be found at https://hyperbrowser.uio.no/assemblygaps. All the data and results are available in the https://hyperbrowser.uio.no/assemblygaps/u/hb-superuser/p/assembly-gaps).

## Author’s contributions

DD performed all the simulations and analysis. DD, CK and GKS participated in the design of the study and drafted the manuscript. DD and BS implemented the analysis functionalities. GKS conceived the idea of the study. All authors read and approved the final manuscript.

## Acknowledgements

Not applicable

## References

1. Consortium, I.H.G.: Finishing the euchromatic sequence of the human genome. Nature 431, 931–945 (2004)

2. An integrated encyclopedia of dna elements in the human genome. Nature 489(7414), 57–74 (2012)

3. Lander, E.S.: Initial impact of the sequencing of the human genome. Nature 470(7333), 187–197 (2011)

4. Treangen, T.J., Salzberg, S.L.: Repetitive dna and next-generation sequencing: computational challenges and solutions. Nature Reviews. Genetics 13(1), 36–46 (2011). doi:10.1038/nrg3117

5. Schneider, V.A., Graves-Lindsay, T., Howe, K., Bouk, N., Chen, H.-C., Kitts, P.A., Murphy, T.D., Pruitt, K.D., Thibaud-Nissen, F., Albracht, D., Fulton, R.S., Kremitzki, M., Magrini, V., Markovic, C., McGrath, S., Steinberg, K.M., Auger, K., Chow, W., Collins, J., Harden, G., Hubbard, T., Pelan, S., Simpson, J.T., Threadgold, G., Torrance, J., Wood, J.M., Clarke, L., Koren, S., Boitano, M., Peluso, P., Li, H., Chin, C.-S., Phillippy, A.M., Durbin, R., Wilson, R.K., Flicek, P., Eichler, E.E., Church, D.M.: Evaluation of grch38 and de novo haploid genome assemblies demonstrates the enduring quality of the reference assembly. Genome Research 27(5), 849–864 (2017). doi:10.1101/gr.213611.116

6. Mouse Genome Assembly GRCm38.p5 Statistics, Genome Reference Consortium. https://www.ncbi.nlm.nih.gov/grc/mouse/data

7. Ebrafish Genome Assembly GRCz11 Statistics, Genome Reference Consortium. https://www.ncbi.nlm.nih.gov/grc/zebrafish/data

8. De, S., Pedersen, B.S., Kechris, K.: The dilemma of choosing the ideal permutation strategy while estimating statistical significance of genome-wide enrichment. Briefings in Bioinformatics 15(6), 919–928 (2014). doi:10.1093/bib/bbt053

9. Ferkingstad, E., Holden, L., Sandve, G.K.: Monte carlo null models for genomic data. Statist. Sci. 30(1), 59–71 (2015). doi:0.1214/14-STS484

10. Quinlan, A.R.: Bedtools: the swiss-army tool for genome feature analysis. Current protocols in bioinformatics / editoral board, Andreas D. Baxevanis … [et al.] 47, 11–121111234 (2014). doi:10.1002/0471250953.bi1112s47

